# Real-time inactivation of airborne SARS-CoV-2 using ultraviolet-C

**DOI:** 10.1101/2022.10.04.510919

**Authors:** Carolina Koutras, Richard L. Wade

## Abstract

COVID-19 is a life-threatening respiratory infection that has had a profound impact on indoor air quality awareness. Ultraviolet-C (UV-C) is a physical disinfection process that triggers microbial inactivation through creating irreversible genetic material damage. An upper room device equipped with germicidal UV-C (UR GUV) was evaluated against airborne SARS-CoV-2 for antimicrobial efficacy using a robust aerosol testing protocol. In 30 minutes, it led to a virucidal efficacy of 99.994 % in a large, room-sized chamber. UR GUV is a promising mitigation strategy for airborne pathogens.

## Background

The coronavirus disease 2019 (COVID-19) pandemic has been tremendously disruptive to human health, societies and health systems globally, as several waves of outbreaks caused more than 6.5 million deaths around the world. Vaccines and public health protective measures have provided considerable protection against serious COVID-19 outcomes, but the emergence of more infectious SARS-CoV-2 variants and post-COVID long term health conditions continue to disrupt the global economy. Moreover, in the United States, the Centers for Disease In addition to COvid-19, influenza still poses a threat to human health (CDC) estimates that contagious respiratory illness caused by influenza viruses that infect the nose, throat, and lungs has resulted in 9-41 million illnesses, 140,000-710,000 hospitalizations and 12,000-52,000 deaths annually between 2010 and 2020.^1-4^ The near absence of viral respiratory infections due to COVID-19 precautions over the past two years is a reminder that these infections need not happen at the rate that we have long assumed was inevitable.

Upper-room germicidal ultraviolet-C (UV-C) irradiation (also known as UR GUV) is an infection prevention tool first indicated for the inactivation of airborne *Mycobacterium tuberculosis* and has subsequently seen broader adoption during COVID-19. UR GUV fixtures create a field of non-ionizing radiation above the heads of indoor room occupants. The irradiation field acts as a disinfection zone leading to the inactivation of airborne microorganisms that move from the breathing zone to the disinfection zone through natural vertical air movement. UV-C light induces damage to the genomes of bacteria, viruses, and other microorganisms by breaking bonds and forming photodimeric lesions in nucleic acids. These lesions prevent both transcription and replication and ultimately lead to microbial inactivation. Despite significant technological advancements in the field, questions remain about the real-world efficacy of UR GUV fixtures as a standard Environmental Protection Agency (EPA) method does not currently exist for evaluating the efficacy of UV-C against airborne pathogens. The goal of this study was to provide a reference for reporting and evaluating the efficacy of UR GUV devices against high concentrations of viable viral aerosols in a patient room-sized chamber using SARS-CoV-2 as a challenge organism.^5-7^

## Methods

Testing was conducted in a sealed 20’x8’x8’ chamber per Biosafety Level 3 (BSL3) standards (Innovative Bioanalysis Laboratory, CA). The overall dimensions of the test chamber provided a displacement volume of approximately 36,245.56 liters of air, meeting and exceeding the standard size for a room according to the EPA and simulates real world testing of disinfection devices against aerosolized microorganisms. A nebulizing port connected to a programmable compressor system was located in the center of the 20 ft wall protruding 24-inches from the wall. At each chamber corner, low-volume mixing fans (approximately 30 cfm each) were positioned at 45-degree angles to ensure homogenous mixing of bioaerosol concentrations when nebulized into the chamber. The room was equipped with four probes for air sampling positioned along the room’s centerline and located 6 ft off the chamber floor. An UR GUV device (Beam, R-Zero Systems) equipped with 48 light emitting diodes (265 nm LEDs) emitting approximately 67 uW/cm2 at 5 feet from the light source was mounted at the center of one of the 8-foot walls, approximately 6.5 ft off the floor (Figure 1). The temperature during all test runs was approximately 21 +/-2 °C, with a relative humidity (RH) of 45%. A Blaustein Atomizing Module (BLAM) with a pre-set PSI and computer-controlled liquid delivery system tested for average particle size distribution and used for bioaerosol generation. The nebulizer was filled with 7.04 × 10^6^ TCID_50_/mL of SARS-CoV-2 (SARS-Related Coronavirus 2, Isolate USA-CA1/2020, NR-52382) in viral suspension media and nebulized at a flow rate of 1mL/min with untreated local atmospheric air. After nebulization, the nebulizer’s remaining viral stock volume was weighed to confirm roughly the same amount was nebulized during each three replicate runs. Bioaerosol sampling occurred at 0, 5, 15, and 30 minutes using four probes for air sampling, each connected to a calibrated Gilian 10i vacuum device and set at a standard flow of 5.02 L/min with a 0.20% tolerance. Before use, the devices were inspected for functionality, and the vacuum system calibration was confirmed using a Gilian Gilibrator-2 NIOSH Primary Standard Air Flow Calibrator. Sample collection volumes were set to 10-minute draws per time point, which allowed for approximately 50 liters of air collection per collection port. The air sampler operated with a removable sealed cassette and was manually removed after each sampling time point. Cassettes had a delicate internal filtration disc (Zefon International) to collect virus samples, which was moistened with a virus suspension media to aid in the collection. At each time point, the device was turned off for sample collection (as applicable). Air sampling collections were set to 10-minute continuous draws. All sample discs were pooled into one collection tube to provide an average across the four sampling locations. After the control runs, all procedures were then repeated with the UR GUV device running in the same manner with corresponding time points and collection rates.

**Figure 1.**
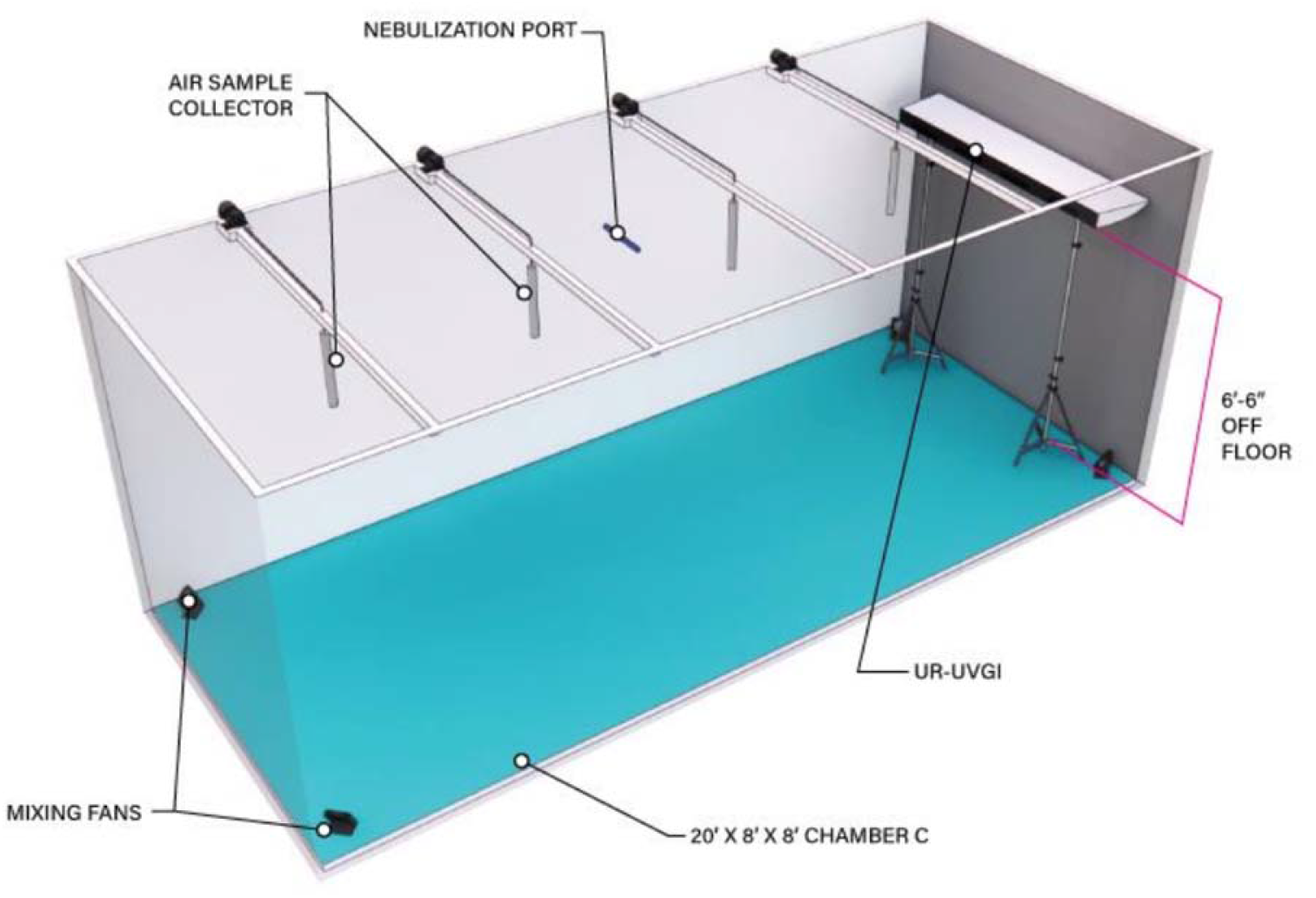
Study Diagram. The UR GUV test device was placed at the center of an 8-foot wall, approximately 6.5 ft off the floor.

After each 30 minutes run, the air filtration system was purged for 30-minutes. Test equipment was cleaned at the end of each day with a 70% alcohol solution. Collection lines were soaked in a bleach bath mixture for 30 minutes then rinsed repeatedly with DI water. The nebulizer and vacuum collection pumps were decontaminated with hydrogen peroxide mixtures.

Following harvest of treated and control discs, the viral suspensions were quantified using the TCID_50_ (Median Tissue Culture infectious Dose) technique. The inoculated cell culture plates were incubated and microscopically scored for the presence/absence of the test virus. The Spearman-Kärber method was used for estimating viral titers. The log10 and percent reductions in viral titer were calculated for UV-C exposed carriers relative to controls.

## Results

Table 1 summarizes the TCID_50_ for all control and UR GUV treated samples, and average TCID_50_ for both test and control conditions at 0, 5, 15 and 30 minutes. As expected, the control showed a natural viability loss of aerosolized SARS-CoV-2 for 30 minutes within the chamber under controlled conditions, however the recovery was meaningful. For three trials against SARS-CoV-2, an initial concentration of 7.04 × 10^6^ TCID_50_/mL was reduced to 4.55 × 10^6^, 4.72 × 10^6^, and 4.25 × 106 TCID_50_/mL, averaging to approximately 4.51 × 10^6^ TCID_50_/mL after 5 minutes of device operation. After 15 minutes, the device decreased collectible SARS-CoV-2 to 1.25 × 10^6^, 9.86 × 10^5^, and 7.47 × 10^5^, averaging to approximately 9.93 × 10^5^ TCID_50_/mL. After 30 minutes, 4.80 × 10^2^, 1.20 × 10^2^, and 1.20 × 10^2^, averaging to 2.40 × 10^2^ TCID_50_/mL.

**Table 1.**
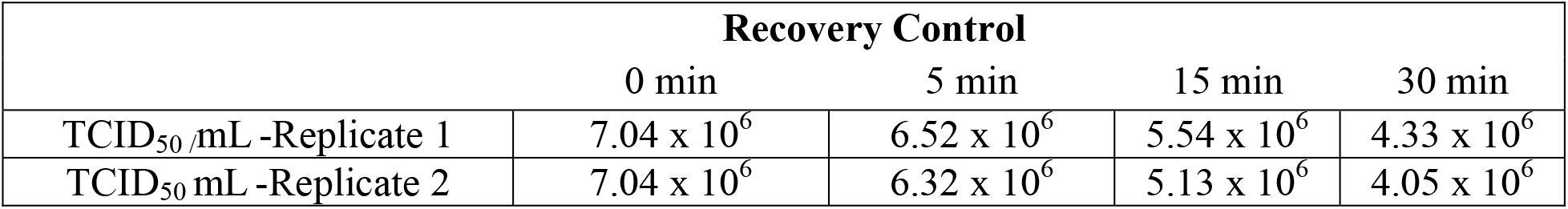
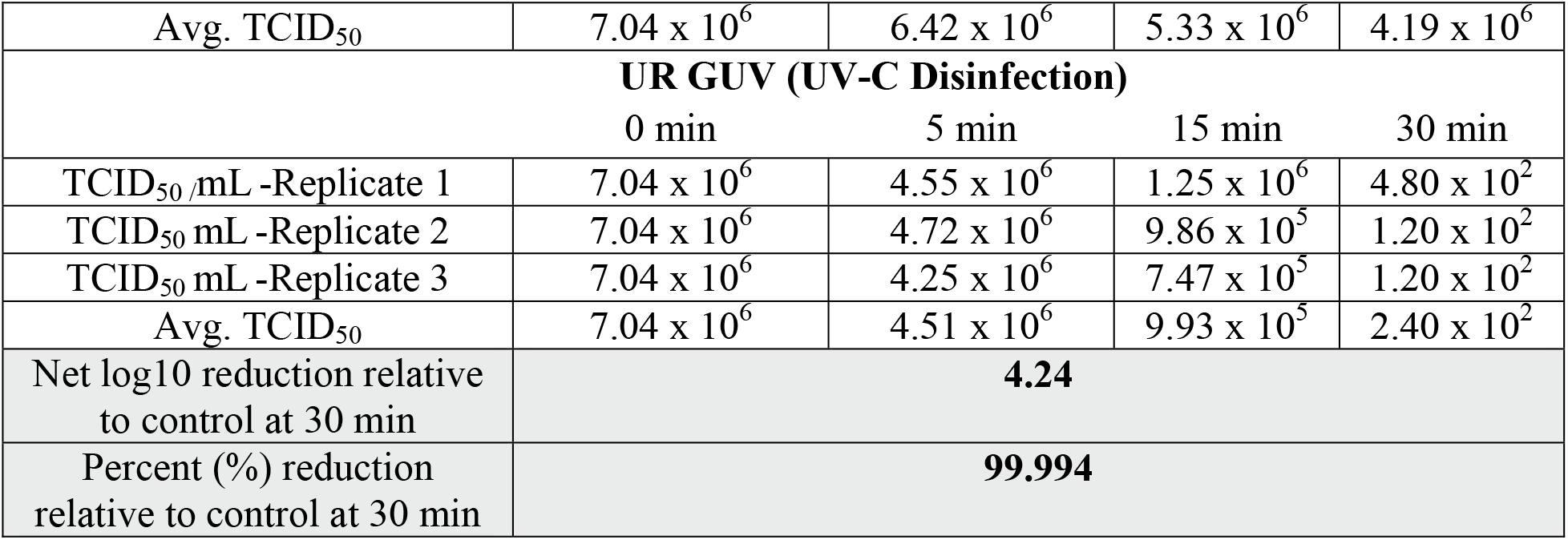
The average TCDI_50_ per time point is shown for both the recovery and test replicates. The average log10 and percent reduction for the test condition relative to control is also provided.

| Recovery Control |  |  |  |  |
| --- | --- | --- | --- | --- |
|  | 0 min | 5 min | 15 min | 30 min |
| TCID<sub>50</sub>/mL -Replicate 1 | $7.04 \times 10^6$ | $6.52 \times 10^6$ | $5.54 \times 10^6$ | $4.33 \times 10^6$ |
| TCID<sub>50</sub> mL -Replicate 2 | $7.04 \times 10^6$ | $6.32 \times 10^6$ | $5.13 \times 10^6$ | $4.05 \times 10^6$ |
| Avg. TCID<sub>50</sub> | 7.04 x 10<sup>6</sup> | 6.42 x 10<sup>6</sup> | 5.33 x 10<sup>6</sup> | 4.19 x 10<sup>6</sup> |
| **UR GUV (UV-C Disinfection)** |  |  |  |  |
|  | 0 min | 5 min | 15 min | 30 min |
| TCID<sub>50</sub>/mL -Replicate 1 | 7.04 x 10<sup>6</sup> | 4.55 x 10<sup>6</sup> | 1.25 x 10<sup>6</sup> | 4.80 x 10<sup>2</sup> |
| TCID<sub>50</sub> mL -Replicate 2 | 7.04 x 10<sup>6</sup> | 4.72 x 10<sup>6</sup> | 9.86 x 10<sup>5</sup> | 1.20 x 10<sup>2</sup> |
| TCID<sub>50</sub> mL -Replicate 3 | 7.04 x 10<sup>6</sup> | 4.25 x 10<sup>6</sup> | 7.47 x 10<sup>5</sup> | 1.20 x 10<sup>2</sup> |
| Avg. TCID<sub>50</sub> | 7.04 x 10<sup>6</sup> | 4.51 x 10<sup>6</sup> | 9.93 x 10<sup>5</sup> | 2.40 x 10<sup>2</sup> |
| Net log10 reduction relative to control at 30 min | **4.24** |  |  |  |
| Percent (%) reduction relative to control at 30 min | **99.994** |  |  |  |

## Discussion

UR GUV is not a novel infection prevention tool, but its adoption in healthcare and community settings has been propelled by COVID-19. In this study, an UR GUV device successfully inactivated 99.994 % (4.24 log10) of SARS-CoV-2 in a large chamber in 30 minutes. Indoor environments can vary greatly, and while a large chamber may not reflect the characteristics of every indoor space exactly, the authors believe that testing UV-C air disinfection technologies in controlled settings is appropriate and important for several reasons. In real world settings outside of rigorous clinical trials, uncontrollable variables such as bioaerosol/bioburden cross-contamination between UV-C treated and adjacent non-treated areas could greatly impact results; pathogens can’t be nebulized safely is most spaces; sampling of bioaerosols require the use of delicate and carefully calibrated equipment, which may be challenging to maintain in the field; and the timing of sampling and disinfection protocols in place may also compromise the results. As such, in the absence of formal regulatory guidance, aerosol testing studies for UR GUV should consider incorporating a detailed description of the volume of the chamber, the microorganism identification/source and concentration, the aerosol generation process, the aerosol sampling process, aerosol dispersion information, the sterility of the chamber, the environmental conditions in the chamber, the irradiance of the device, and contact time needed to demonstrate effectiveness. In conclusion, the tested device was proven highly effective against airborne SARS-CoV-2 using a robust aerosol testing protocol that can be referenced by manufacturers and infection preventionists to evaluate the efficacy of other UR GUV technologies and de-risk implementation.

## Acknowledgements

The authors would like to thank Josh Eddy and Innovative Bioanalysis Laboratories (Costa Mesa, CA) for their commitment to this project.

